# Neural correlates of cue-induced changes in decision-making distinguish subjects with gambling disorder from healthy controls

**DOI:** 10.1101/498725

**Authors:** Alexander Genauck, Caroline Matthis, Milan Andrejevic, Lukas Ballon, Francesca Chiarello, Katharina Duecker, Andreas Heinz, Norbert Kathmann, Nina Romanczuk-Seiferth

**Author notes:** **Corresponding Author**: Alexander Genauck, Department of Psychiatry and Psychotherapy, Charité - Universitätsmedizin Berlin, Charitéplatz 1, 10117 Berlin, Germany. **Funding Sources**: This study was funded by a research grant by the Senatsverwaltung für Gesundheit, Pflege und Gleichstellung, Berlin. A.G. was funded by Deutsche Forschungsgemeinschaft (DFG) HE2597/15-1, HE2597/15-2, and DFG Graduiertenkolleg 1519 “Sensory Computation in Neural Systems”. **Authors’ contribution**: AG designed the experiment, collected the data, analyzed the data, wrote the manuscript. CM reviewed the machine-learning algorithm and revised the manuscript. MA collected data, revised manuscript. AH revised the manuscript, oversaw manuscript drafting. NK revised the manuscript, advised first author. FC analyzed data, revised manuscript. KD collected data, analyzed data, revised manuscript. NRS designed and supervised study and experiment, oversaw manuscript drafting and data analysis. **Co-Authors’ email addresses**: Caroline Matthis; Andreas Heinz; Norbert Kathmann; Nina Romanczuk-Seiferth; Francesca Chiarello; Katharina Duecker; Milan Andrejevic; Lukas Ballon. **Remarks**: To ensure a more convienient reviewing process, we positioned figures and tables at their destined postion.

## Abstract

**Background:** Just as substance use disorders (SUD), gambling disorder (GD) is characterized by an increase in cue-dependent decision-making (Pavlovian-to-instrumental transfer, PIT). PIT, as studied in SUDs and healthy subjects, is associated with altered communication between Nucleus Accumbens (NAcc), amygdala, and orbitofrontal cortex (OFC). However, these neural differences are poorly understood. For example, it is unclear whether they are due to the physiological effects of substance abuse, or rather related to learning processes and/or other etiological factors like innate traits associated with addiction. We have thus investigated whether network activation patterns during a PIT task are also altered in GD, an addictive disorder not involving substance abuse. We have specifically studied which *neural* PIT patterns were best at distinguishing GD from healthy control (HC) subjects, all to improve our understanding of the neural signatures of GD and of addiction-related PIT in general.

**Methods:** 30 GD and 30 HC subjects completed an affective decision-making task in a functional magnetic resonance imaging (fMRI) scanner. Gambling associated and other emotional cues were shown in the background during the task, allowing us to record multivariate neural PIT signatures focusing on a network of NAcc, amygdala and OFC. We built and tested a classifier based on these multivariate neural PIT signatures using cross-validated elastic net regression.

**Results and Discussion:** As expected, GD subjects showed stronger PIT than HC subjects because they showed stronger increase in gamble acceptance when gambling cues were presented in the background. Classification based on neural PIT signatures yielded a significant AUC-ROC (0.70, p = 0.013). When inspecting the features of the classifier, we observed that GD showed stronger PIT-related functional connectivity between nucleus accumbens (NAcc) and amygdala elicited by gambling background cues, as well as between amygdala and orbito-frontal cortex (OFC) elicited by negative and positive cues.

**Conclusion:** We propose that GD and HC subjects are distinguishable by PIT-related *neural* signatures including amygdala-NAcc-OFC functional connectivity. Our findings suggest that neural PIT alterations in addictive disorders might not depend on the physiological effect of a substance of abuse, but on related learning processes or even innate neural traits, also found in behavioral addictions.

## INTRODUCTION

Gambling disorder (GD) has been classified as an addiction alongside substance-use disorders (SUDs), such as alcohol or cocaine dependence (American Psychiatric Association, American Psychiatric Association, & DSM-5 Task Force, 2013). This new classification was indicated because GD and SUDs share the same core symptoms (including craving, withdrawal, tolerance) and both GD and SUDs show similar neuro-behavioral signatures (Clark, 2014; Clark et al., 2013; Leeman & Potenza, 2012; N. M. Petry et al., 2014; Romanczuk-Seiferth, van den Brink, & Goudriaan, 2014).

For instance, just like patients suffering from SUDs, GD subjects show increased neural activity elicited by addiction-related stimuli (i.e. “cues”) and a reduced neural response towards stimuli signaling natural rewards (Carter & Tiffany, 1999; Crockford, Goodyear, Edwards, Quickfall, & el-Guebaly, 2005; Goudriaan, de Ruiter, van den Brink, Oosterlaan, & Veltman, 2010; Potenza et al., 2003; Rømer Thomsen, Fjorback, Møller, & Lou, 2014). In addiction, a cue can be any formerly neutral stimulus that has been repeatedly paired with the effects of the addictive behavior (Mucha, Geier, Stuhlinger, & Mundle, 2000; Potenza et al., 2003). The effect of increased responsivity towards addiction-related cues is termed cue reactivity and is pivotal in explaining a range of behaviors related to addictive disorders, such as arousal, attentional bias, craving, and relapse (A. Beck et al., 2012; Carter & Tiffany, 1999; Field, Munafò, & Franken, 2009; Goudriaan et al., 2010; Heinz et al., 2003; Leyton & Vezina, 2012, 2013; Schacht, Anton, & Myrick, 2013; Vezina & Leyton, 2009; Wölfling et al., 2011). Besides cue reactivity, and just like in SUDs, GD subjects display impaired value-based decision-making. For example, GD subjects show increased risk taking, higher discounting of delayed rewards (delay discounting) and reduced loss aversion (Clark et al., 2013; Dixon, Marley, & Jacobs, 2003; Genauck et al., 2017; Glimcher & Rustichini, 2004; Lorains et al., 2014; MacKillop et al., 2011; Madden, Petry, & Johnson, 2009; N. M. Petry, 2012; Platt & Huettel, 2008; Romanczuk-Seiferth et al., 2014; Wiehler & Peters, 2015). Impaired value-based decision-making in addiction may partly be explained, or even further exacerbated by, cues that modulate decision-making processes. The modulating influence of conditioned cues on instrumental behavior (i.e. increasing the vigor with which a behavior is displayed or increasing the likelihood of choosing a certain option) has been termed Pavlovian-to-instrumental transfer (PIT) (Cartoni, Balleine, & Baldassarre, 2016; De Tommaso, Mastropasqua, & Turatto, 2018; Schulreich, Gerhardt, & Heekeren, 2016; Talmi, Seymour, Dayan, & Dolan, 2008). PIT is one of the key effects deepening our understanding of cue-controlled behaviors (Dickinson & Balleine, 1994; Dickinson Anthony & Balleine Bernard, 2002; Holmes, Marchand, & Coutureau, 2010; Niv, Daw, Joel, & Dayan, 2007). Interestingly, PIT effects can persist even when the outcome of the instrumental behavior has been devalued (De Tommaso et al., 2018), and a stronger PIT has been associated with heightened impulsivity (Garofalo & Robbins, 2017) and with reduced model-based behavior (Sebold et al., 2016). This is why PIT has gained considerable attention in the field of addiction research. Patients suffering from SUDs have displayed increased PIT (Corbit & Janak, 2016; Corbit, Janak, & Balleine, 2007; Corbit, Nie, & Janak, 2012; Garbusow et al., 2016; Schad et al., 2018), however not in all studies (Hogarth & Chase, 2012). Furthermore, the effect seems to scale with the probability of relapse (Garbusow et al., 2016).

Investigating PIT in GD is of particular importance to understanding addictive disorders in general, because GD is an addictive disorder independent of any neurotropic substance of abuse. The study of PIT in GD therefore helps us distinguish whether PIT effects seen in SUDs are a physiological result caused by the abused substance, or rather by addiction-related learning (Heinz, 2017, p. 113ff.), or even by innate traits putatively associated with developing and maintaining an addiction (Barker, Torregrossa, & Taylor, 2012).

So far only a small number of studies have investigated PIT in GD subjects. It has been observed in GD subjects that delay discounting is increased under the influence of high-craving gambling cues vs. low-craving gambling cues (Dixon, Jacobs, & Sanders, 2006; Miedl, Büchel, & Peters, 2014). GD subjects also have shown to be more strongly influenced by gambling cues in a response inhibition task than HC subjects (van Holst, van Holstein, van den Brink, Veltman, & Goudriaan, 2012). To investigate PIT in GD, Genauck et al. (under review) used a mixed-gambles task, i.e. a task where participants have to decide whether they want to take gambles that entail both possible gains and losses. They coupled the task with emotional and gambling-related cues (affective mixed-gambles task) to estimate subject-specific behavioral PIT parameters with regards to loss aversion. The authors found that behavioral PIT parameters lend themselves to classify subjects into HC vs. GD subjects. The most successful model to separate GD subjects from HC subjects was the one explaining the shift in general gamble acceptance by the influence of different cue categories, while loss aversion and loss-aversion specific PIT did not improve the distinction between GD from HC. In the present study, subjects performed a very similar affective mixed-gambles task in a functional magnetic resonance imaging (fMRI) scanner. Genauck et al. (under review) successfully used the behavioral data of the present study as an independent sample to validate their classifier.

We mentioned studies which suggest that GD is associated with increased PIT, despite the disorder being independent of any substance of abuse. However, it is unclear if there are also *neural* PIT signatures associated with cue-induced decision-making which distinguish GD from HC subjects, just like there are between SUD and HC subjects. If there are neural PIT signatures associated with GD then this would be additional evidence for functional brain changes related to addictive disorders independent of a substance of abuse (Goudriaan, de Ruiter, van den Brink, Oosterlaan, & Veltman, 2010; Koehler et al., 2013; Romanczuk-Seiferth, Koehler, Dreesen, Wüstenberg, & Heinz, 2015; Sescousse, Barbalat, Domenech, & Dreher, 2013; van Holst, van der Meer, et al., 2012; van Holst, van Holstein, van den Brink, Veltman, & Goudriaan, 2012). Our study is the first to investigate functional brain changes in GD compared to HC related to cue-induced changes in value-based decision-making. We expected that neural PIT signatures derived from SUD studies should underlie behavioral PIT increase also in GD, and thus lend themselves to distinguish GD from HC subjects.

At the neural level, PIT depends on the functions of amygdala and the ventral striatum (VS) (Corbit & Balleine, 2005; Corbit, Muir, & Balleine, 2001; de Borchgrave, Rawlins, Dickinson, & Balleine, 2002; Hall, Parkinson, Connor, Dickinson, & Everitt, 2001; Prévost, Liljeholm, Tyszka, & O’Doherty, 2012; Talmi et al., 2008). The VS denotes the ventral parts of caudate and putamen in humans and it is often used interchangeably with the nucleus accumbens (NAcc) region. For example, a study found evidence for the contribution of the ventral putamen in PIT (Bray, Rangel, Shimojo, Balleine, & O’Doherty, 2008). In addition, Garbusow et al. (2016) distinguished alcohol dependent relapsers from abstainers using a nucleus accumbens (NAcc) PIT signal, reaching an accuracy of 71% in leave-one-out cross-validation. Note that cue reactivity, which PIT arguably is based upon, is also associated with altered activity of amygdala and NAcc in addictive disorders (Kühn & Gallinat, 2011; Schacht et al., 2013).

In addition to possible activity differences in limbic regions being associated with PIT, recent literature suggests that functional NAcc-amygdala connectivity plays a role in decision-making changes due to emotional cues (Charpentier, Martino, Sim, Sharot, & Roiser, 2015). Other authors have argued that Pavlovian influence on instrumental behavior require the modulation of ongoing processes in the striatum by the amygdala (Guitart-Masip, Talmi, & Dolan, 2010), (Cardinal, Parkinson, Hall, & Everitt, 2002). Bi-directional NAcc-amygdala connectivity could thus be enhanced in GD subjects during presentation of addiction-relevant cues. Holmes et al. (2010) and Cardinal et al. (2002) further suggest a contribution of the orbital frontal cortex in integrating information about Pavlovian and instrumental processes, together with the striatum and amygdala. The ANDREA (affective neuroscience of decision through reward-based evaluation of alternatives) model makes similar predictions when explaining transient changes in gamble acceptance in decision-making tasks (Litt, Eliasmith, & Thagard, 2008) (**Fig. 2**). In particular, this model suggests that the evaluation of a gamble involving possible gains and losses leads to a subjective value signal in the OFC. Amygdala inputs to OFC can modulate those subjective value representations when e.g. gambling cues are shown in the background. Since GD subjects show stronger behavioral PIT effects related to gambling cues, this could mean that gambling cues increase the subjective gamble value represented in OFC via amygdala projections. We thus expected that stronger gambling-cue PIT-related functional connectivity from amygdala to OFC should help distinguish GD from HC.

In summary, we hypothesized that a neural PIT signature made up of several PIT-related fMRI contrasts could distinguish GD from HC subjects. We therefore compiled a feature vector comprised of cue reactivity and PIT-related contrasts in amygdala and NAcc, and of functional connectivity parameters in a network of NAcc, amygdala and OFC. Hence the feature vector represented each subject’s neural PIT signature, in the form of multiple functional magnetic resonance imaging (fMRI) aggregates (Seo et al., 2018, 2015; Whelan et al., 2014). We used all subjects’ neural PIT signatures to estimate a classifier which would distinguish GD from HC subjects. We expected that PIT-related predictors would be found among the most important ones followed by the cue-reactivity predictors. Using cross-validation we assessed the generalizability of this classifier to new samples. Classifying GD and HC subjects using multivariate patterns aims to bring us closer to a clinically relevant characterization of the neural disturbances related to GD, especially when there are many relevant variables involved (Ahn, Ramesh, Moeller, & Vassileva, 2016; Ahn et al., 2016; Cerasa et al., 2018; Guggenmos et al., 2018; Yarkoni & Westfall, 2017). To our knowledge, our study is the first one to use fMRI-based classification for investigating GD and its neural basis of increased PIT.

## METHODS AND MATERIALS

### Sample

The GD group consisted of subjects who where active gamblers, while the HC group consisted of subjects that had none or little experience in gambling and did not display problematic gambling. We recruited GD subjects via eBay classifieds, and notices in Berlin casinos and gambling halls. GD subjects were diagnosed using the German short questionnaire for gambling behavior (KFG) (cutoff ≥ 16) (J. Petry & Baulig, 1996). The KFG classifies subjects according to DSM-IV criteria for pathological gambling. However, in the following we use the DSM-5 term “gambling disorder” interchangeably, because the criteria largely overlap (Rodríguez-Testal, Senín-Calderón, & Perona-Garcelán, 2014). Any known history of a neurological disorder or a current psychological disorder (except tobacco dependence) as assessed by the Screening of the Structured Clinical Interview for DSM-IV Axis I Disorders (SCID-I) (First, Spitzer, Gibbon, & Williams, 2002) led to exclusion from the study. For further information on administered questionnaires, see Supplements. There were 13 subject dropouts due to technical errors, positive drug screenings, incidental cerebral anatomical findings or MRI contraindications. We dropped five more subjects to improve the matching of the groups on covariates of no interest (age, smoking severity, education, and see below). The final sample consisted of 30 GD and 30 HC subjects (Tab. 1). GD subjects differed in gambling habits to HC mainly in frequency of playing slot machines (GD: once a week or more, HC: never or less than once a week, p < 0.001) and in casinos (GD: around once a week, HC: never or less than once a week, p < 0.001). GD and HC were matched on relevant variables (net personal income, age, alcohol use), except for years in school (primary and secondary). We thus tested for stability of our classifier by adjusting for years in school.

**Table 1:** Sample characteristics, means and p-values calculated by two-sided permutation test.

| variable | HC (30) | se | PG (30) | se | pooled se | p perm test |
| --- | --- | --- | --- | --- | --- | --- |
| years in school | 10.87 | 0.19 | 10.13 | 0.24 | 0.21 | 0.031 |
| vocational school | 2.73 | 0.29 | 2.07 | 0.25 | 0.27 | 0.108 |
| net personal income | 1028.61 | 92.27 | 1105.89 | 138.93 | 115.6 | 0.667 |
| personal debt | 8500 | 3396.88 | 24000 | 9590.36 | 6493.62 | 0.097 |
| Fagerström | 1.97 | 0.43 | 3.03 | 0.51 | 0.47 | 0.138 |
| age | 35.37 | 1.66 | 37.37 | 2.01 | 1.84 | 0.459 |
| AUDIT | 4.8 | 0.59 | 4.87 | 1.05 | 0.82 | 1 |
| BDI-II | 5.1 | 1.03 | 11.57 | 1.72 | 1.38 | 0.002 |
| SOGS | 1.73 | 0.47 | 8.8 | 0.67 | 0.57 | <0.001 |
| KFG | 2.37 | 0.74 | 35 | 1.64 | 1.19 | <0.001 |
| BIS-15 | 31.8 | 0.99 | 36.33 | 1.08 | 1.03 | 0.004 |
| GBQ persistence | 1.96 | 0.2 | 3.28 | 0.19 | 0.2 | <0.001 |
| GBQ illusions | 2.41 | 0.24 | 3.73 | 0.22 | 0.23 | <0.001 |
| ratio female | 0.20 | - | 0.20 | - | - | 1.000 |
| ratio unemployed | 0.17 | - | 0.20 | - | - | 1.000 |
| ratio smokers | 0.60 | - | 0.77 | - | - | 0.262 |
| ratio right-handed | 0.97 | - | 0.84 | - | - | 0.204 |

\*chi-square test used; se: bootstrapped standard errors; years in school: years in primary and secondary school; vocational school is vocational school and university; Fagerström: smoking severity (Heatherton, Kozlowski, Frecker, & Fagerström, 1991); AUDIT: alcohol use disorders identification test (Dybek et al., 2006); BDI II: Beck's Depression Inventory (A. T. Beck, Steer, & Brown, 1996), SOGS: South Oaks Gambling Screen according to DSM-III (Lesieur & Blume, 1987); KFG: Kurzfragebogen zum Glückspielverhalten, Short Questionnaire Pathological Gambling, German diagnostic tool and severity measure based on the DSM-IV (J. Petry & Baulig, 1996); BIS-15: short version of the Barratt Impulsiveness Scale for impulsivity (Meule, Vögele, & Kübler, 2011; Patton, Stanford, & Barratt, 1995); GBQ persistence and GBQ illusions: from the Gamblers' Beliefs Questionnaire (Steenbergh, Meyers, May, & Whelan, 2002)

### Procedure and data acquisition

Before scanning, all subjects underwent urine drug testing to exclude any influence of cannabis, amphetamines, cocaine, methamphetamines, opiates, or benzodiazepines. They then were instructed on the task and completed the PIT task in a 3-Tesla SIEMENS Trio MRI (2 runs of about 23 minutes). EPI scans were acquired, as well as structural MRI. For further details on MRI sequences see Supplements.

### Affective mixed-gambles task

We were inspired by established mixed-gambles decision-making tasks (Genauck et al., 2017; Tom, Fox, Trepel, & Poldrack, 2007) and mixed-gambles decision-making tasks with the influence of affective cues (Charpentier et al., 2015; Genauck et al., under review). As affective cues, four sets of images were assembled: 1) 67 gambling images, showing a variety of gambling scenes, and paraphernalia (*gambling cues*); 2) 31 images showing negative consequences of gambling (*negative cues*); 3) 31 images showing positive effects of abstinence from gambling (*positive cues*); 4) 24 neutral IAPS images (*neutral cues*). The cues of all categories were presented in random order and each gambling cue only appeared once. For negative, positive, and neutral cue categories, we randomly drew images from each pool until we had presented 45 images of each category and each image at least once. Hence, we ran 202 trials in each subject. Subjects were each given 20€ for wagering. Every trial began with the presentation of one of the cues described above that subjects were instructed to remember for a paid recognition task after the experiment. After 4s (jittered), a mixed gamble, involving a possible gain and a possible loss, with probability P = 0.5 each, was superimposed on the cue. After another 4s (jittered) of decision time, we asked subjects to indicate how willing they were to accept the gamble by button press. This way we kept decision and motor processes apart. Subjects had to choose how willing they were to accept the gamble on a 4-point Likert-scale to ensure task engagement (Tom et al., 2007) (**Fig. 1**). Gambles were created by randomly drawing with replacement from a matrix with possible gambles consisting of 12 levels of gains (14, 16,…, 36) and 12 levels of losses (−7, -8,…, -18) (**Fig. 1**) (Genauck et al., 2017; Tom et al., 2007; Tversky & Kahneman, 1992). In every subject, we stratified gambles according to mean and variance of gain, loss, gamble variance, and Euclidean distance from gamble matrix (*ed*, i.e. gamble difficulty). We informed subjects that after completing the experiment five of their gamble decisions with ratings of “somewhat yes” or “yes” would be randomly chosen and played for real money.

**Figure 1:**
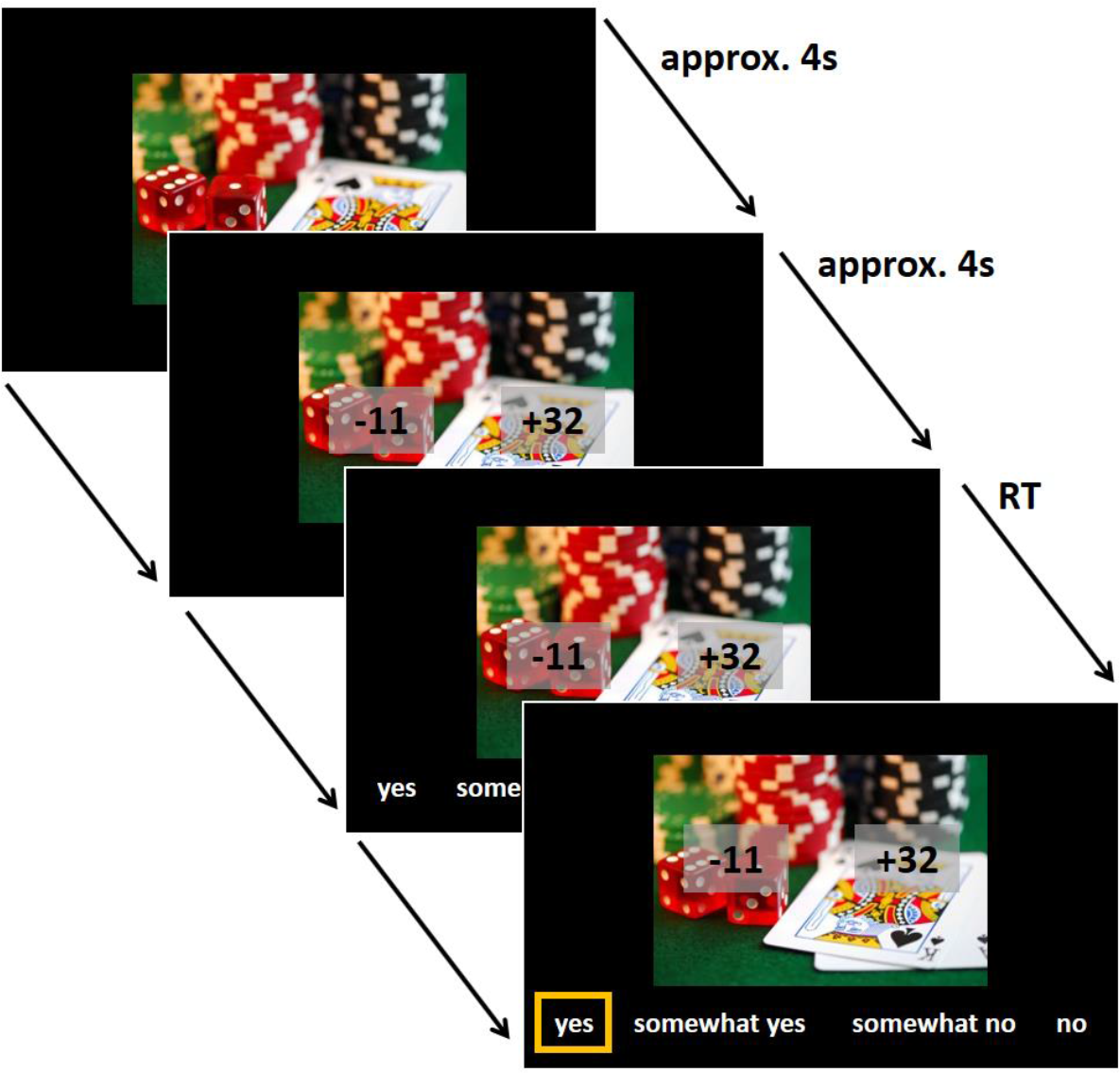
The affective mixed-gambles task. One trial is depicted. Subjects first saw a fixation cross with variable inter-trial-interval (ITI). Then a cue with randomly chosen affective content (67 drawn from 67 gambling related, 45 drawn from 31 with positive consequences of abstinence, 45 drawn from 31 with negative consequences of gambling, 45 drawn from 24 neutral images) was presented for about 4s. Subjects were instructed to remember the cue for a paid recognition task after all trials. Then a gamble involving a possible gain and a possible loss was superimposed on the cue (e.g. -11 and +32). Subjects were instructed to shift their attention at this point to the proposed gamble and evaluate it. Position of gain and loss was counterbalanced (left/right). Gain was indicated by a ‘+’ sign and loss by a ‘−’ sign. After 4s (jittered) subjects were asked to make a choice between four levels of acceptance (yes, somewhat yes, somewhat no, no; here translated from German version which used “ja, eher ja, eher nein, nein”). Direction of options (from left to right or vice versa) and side of gain amount was random. Directly after decision, the ITI started. If subjects failed to make a decision within 4s, ITI started and trial was counted as missing. RT: reaction time.

### Cue ratings

After the affective mixed-gambles task, subjects rated all cues using the Self-Assessment Manikin (SAM) assessment technique (valence, arousal, dominance) (Bradley & Lang, 1994) and additional visual analogue scales. Additional questions were: 1) “How strongly does this image trigger craving for gambling?”; 2) “How appropriately does this image represent one or more gambles?”; 3) “How appropriately does this image represent possible negative effects of gambling?”; 4) “How appropriately does this image represent possible positive effects of gambling abstinence?”. All cue ratings were z-standardized within subject. Cue ratings were analyzed one-by-one using linear mixed-effects regression, using lmer from the lme4 package in R (Bates, Mächler, Bolker, & Walker, 2015), where cue category (and, in the respective models, clinical group) denoted the fixed effects and subjects and cues denoted the sources of random effects.

### Behavioral data

#### Describing the behavioral choice data

Our main focus in this study was to investigate whether there are neural PIT signatures that lend themselves to distinguishing GD from HC subjects. Therefore, we present here merely explorative analyses to describe the behavioral choice data, especially with respect to shifts in general gamble acceptance, while our main focus are the fMRI data. We modeled the choice data within each subject’s behavioral data by submitting dichotomized choices (somewhat no & no: 0; somewhat yes & yes: 1) into a logistic regression. We dichotomized choices to increase the precision when estimating behavioral parameters, in line with previous studies (Barkley-Levenson, Van Leijenhorst, & Galván, 2013; Genauck et al., under review, 2017; Tom et al., 2007). Predictors were centralized values of gain, centralized absolute values of loss, euclidean distance (*ed*) from gamble matrix as indicator of gamble difficulty (Tom et al., 2007) (all down-sampled from twelve to four steps) (Genauck et al., under review, 2017; Tom et al., 2007), and cue category (**c**). We defined the gamble value (*Q*), according to a modeling of the affective mixed-gambles task that has been observed to be successful in distinguishing GD from HC subjects (Genauck et al., under review) and which is also useful for modeling the fMRI data (Genauck et al., 2017; Tom et al., 2007). According to these findings we focus on an additive change in gamble acceptance depending on the cues presented in the background. In particular, we focus here on “the loss aversion with gamble difficulty (*ed*) as additional predictor plus the modulation of acceptance by category” (**laec**) model:

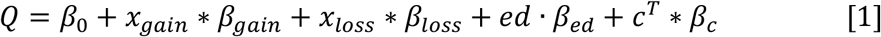

Here *C^T^* is a transposed column vector, denoting the dummy code of the cue’s category on any given trial and *β*_*c*_ is a column vector holding the regression weights describing the shift in gamble value with respect to the cue category. Hence, *C^T^* * *β*_*c*_ is a scalar product describing the additive affective of cue category. For between-group analyses (HC vs. GD) of acceptance rate, and of the cue category effects, we fit the logistic regression based on Eq. [1] with…

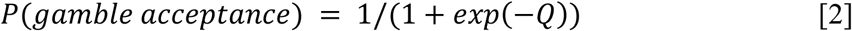

within a generalized linear mixed-effects model, using glmer from the lme4 package in R (Bates et al., 2015). Here, gain, loss, *ed*, cue category denoted the fixed effects and subjects and cues denoted the sources of random effects. To test if the groups differed in the parameters of the **laec** model, we expanded the model by an additional fixed effect of group modulating the effect of gain, loss, *ed*, and cue category (**laecg**). Statistical testing of the model comparison was performed using χ^2^-square difference tests, as well as the comparison of Akaike and Bayesian information criterion (AIC, BIC). For statistical tests of single parameters in the **laecg** model, we used Wald z-test as implemented in lme4. For more analyses of the behavioral data, please see **Supplements**.

### FMRI data

#### Preprocessing and single-subject model of fMRI data

Imaging analyses were performed in SPM12 running on Matlab (R2014a). Please see Supplements for description of preprocessing of MRI data. We modeled the preprocessed fMRI single-subject data using a first onset regressor, which was a boxcar function denoting moments of cue presentation vs. none presentation (“cue on”, 1 vs. 0). This onset regressor had three parametric modulators, denoting gambling/negative/positive “cue on” vs. neutral “cue on” (1 vs. -1). A second onset regressor modeled the time when gamble presentation was on (1 vs. 0). This phase was modeled according to the **laec** model and according to modeling PIT neurally (Garbusow et al., 2016; Schad et al., 2018). Modeling PIT neurally simply entails introducing interaction terms of cue reactivity regressors (e.g. “gambling cue on”) with gamble acceptance regressor (“gamble accepted vs. not accepted”). We therefore had three linearly scaled mean-centered parametric modulators (gain, loss, *ed*, down-sampled from twelve to four steps). We added another parametric modulator denoting acceptance of gamble vs. non-acceptance (1 vs. -1), and three parametric modulators denoting PIT for the three cue categories. For example, the PIT regressor “acceptXgambling” regressor modeled “acceptance during gambling cues vs. not accepting during gambling cues” vs. “accepting during neutral cues vs. not accepting during neutral cues”, i.e. (1 vs. -1) vs. (1 vs. -1). A third onset regressor modeled the onset of the motor response options. We thus had three onset regressors, where the first had three, the second had seven and the third had zero parametric modulators. We modeled missing trials with a missing regressor (1 vs. 0), with duration set at length of trial. Parametric regressors for each onset regressor were serially orthogonalized. Regressors were convolved with the canonical hemodynamic response function, downsampled to match the number of EPI scans and entered into a GLM.

#### Extracting fMRI features for classifier building

We were interested whether PIT fMRI contrasts from a number of brain regions (regions of interest, ROIs) could predict if a subject belongs to the HC or the GD group. We hence extracted the mean activity for cue reactivity (gambling, negative, positive) and for the PIT contrasts (acceptXgambling, acceptXnegative, acceptXpositive) using the within-subject mean from the ROIs NAcc R/L and amygdala R/L. NAcc and amygdala ROIs were taken from the Neuromorphometrics SPM12 brain atlas.

To keep in line with accounts of PIT depending on NAcc-Amy connectivity (Charpentier et al., 2015; Guitart-Masip et al., 2010) and on amygdala-OFC connectivity (Holmes et al., 2010; Litt et al., 2008) (**Fig. 2**), we also extracted functional connectivity (generalized psycho-physiological interaction, gPPI) (McLaren, Ries, Xu, & Johnson, 2012) for the PIT contrasts. We used the seeds amygdala R/L and NAcc R/L. For the seeds amygdala R/L we extracted the mean from target ROIs OFC R/L (4 subregions on either side), and from target ROIs NAcc R/L. For the seeds NAcc R/L, we extracted from the target ROIs Amy R/L. Information from left medial OFC was not available due to signal loss in that region. Note, that at this point we had a vector representing the subjects specific neural PIT pattern. We z-standardized this vector for each subject. We then reduced the dimensionality of this vector for each subject by computing means (For cue reactivity: mean between respective left and right ROI;For functional connectivity: mean connectivity value between respective left and right ROI with respect to each PIT contrast, e.g. for the connectivity from NAcc to posterior OFC with respect to the PIT contrast “acceptXgambling” the mean of connectivity values from R NAcc to R posterior OFC, from L NAcc to R posterior OFC, from R NAcc to L posterior OFC, and from R NAcc to L posterior OFC).

**Figure 2:**
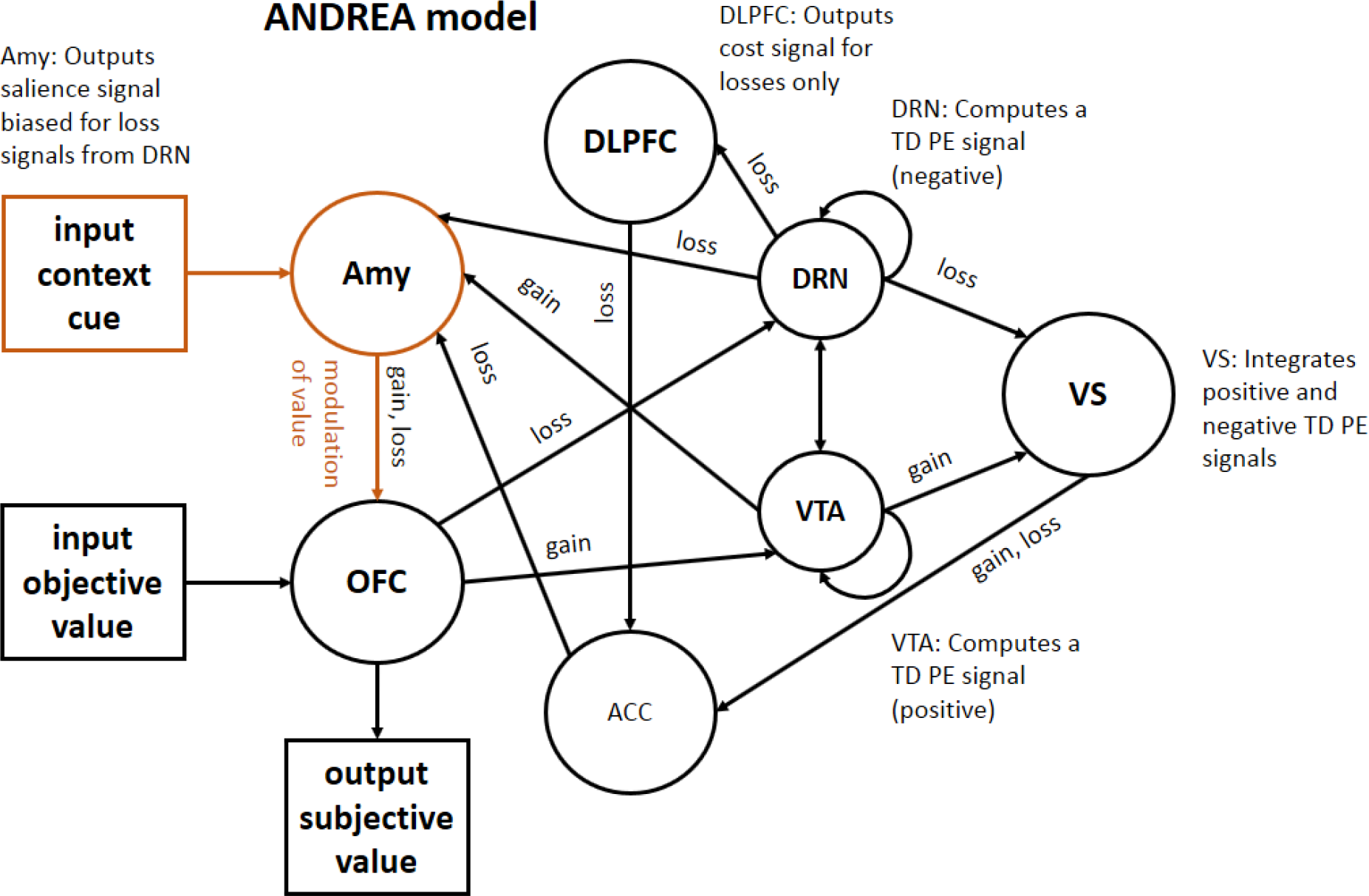
The ANDREA model. The model describes how loss aversion may arise in the brain during a mixed-gambles task and in addition the model makes a specific prediction how contextual cues can influence the subjective representation of gain and loss (this part of the model is highlighted in red). Namely, the amygdala is encoding and forwarding the value signal of the contextual cue, thereby modulating the subjective value representation in OFC (Litt et al., 2008). GD subjects should show a stronger functional connectivity from amygdala to OFC with respect to accepting gambles during presentation of e.g. gambling cues because this would increase the value of the gamble stored in OFC into positive direction and thus increase the likelihood of gamble acceptance.

#### Building the classifier based on fMRI data

The *neural* PIT vectors per subject were stacked into a data set. Since HC and GD were not perfectly matched on years of education, we added this variable to the data set, which was then submitted to generalized elastic net regression, with group as dependent variable. Elastic net regression is well suited for cases where there are few observations and many predictor variables that may contain groups of correlated variables (Zou & Hastie, 2005). Using tuning of its two hyper-parameters (Zou & Hastie, 2005) it is also well suited to produce models that do not over-fit but generalize well to new data, especially when using cross-validation for tuning (Arlot & Celisse, 2010; Bratu, Muresan, & Potolea, 2008; Varma & Simon, 2006). The algorithm tuned for optimal generalization performance on out-of-sample data using the area under the receiver-operating curve, AUC-ROC, (Ahn et al., 2016; Ahn & Vassileva, 2016; Whelan et al., 2014; Zacharaki et al., 2009). AUC-ROC ranges from 0.5 (chance) to 1 (perfect sensitivity and specificity) (Provost, Fawcett, & Kohavi, 1998).

We assessed the generalizability of the above algorithm 1000 times via 10-fold cross-validation (Arlot & Celisse, 2010) which yielded a distribution of classifiers and thus of AUC-ROC’s. Note that the cross-validation to estimate generalizability led to the cross-validations used in the elastic net regression to become *nested* (Arlot & Celisse, 2010; Bratu et al., 2008; Varma & Simon, 2006; Whelan et al., 2014). For a graphical illustration of the algorithm with cross-validation to estimate the generalization performance, see **Fig. 3**. The data and R Code can be found here: https://github.com/pransito/PIT_GD_MRI_release. We computed the mean of the obtained AUC-ROC’s and estimated its p-value by performing the exact same 1000 CV rounds but each time with only “years of education” as predictor (null-classifier). We then subtracted the AUC-ROC’s of the null-classifiers one-by-one from the 1000 AUC-ROC’s of the full classifiers. This yielded a bootstrapped distribution of classification improvement (i.e., improvement of AUC-ROC due to using the full classifier instead of the null-classifier). We tested this distribution against the value of classification improvement under the null-hypothesis (i.e. zero improvement) to obtain a p-value of significance of classification improvement.

After assessing the generalizability of the model by cross-validation, we then fit the model to the entire data set (no splitting in training and test data) in order to build the final interpretable and reportable classifier. Since the modelling is probabilistic, we repeated this 1000 times. We plotted the ensuing distribution of regression weight vectors as per-parameter means with 95% percentile bounds.

**Figure 3:**
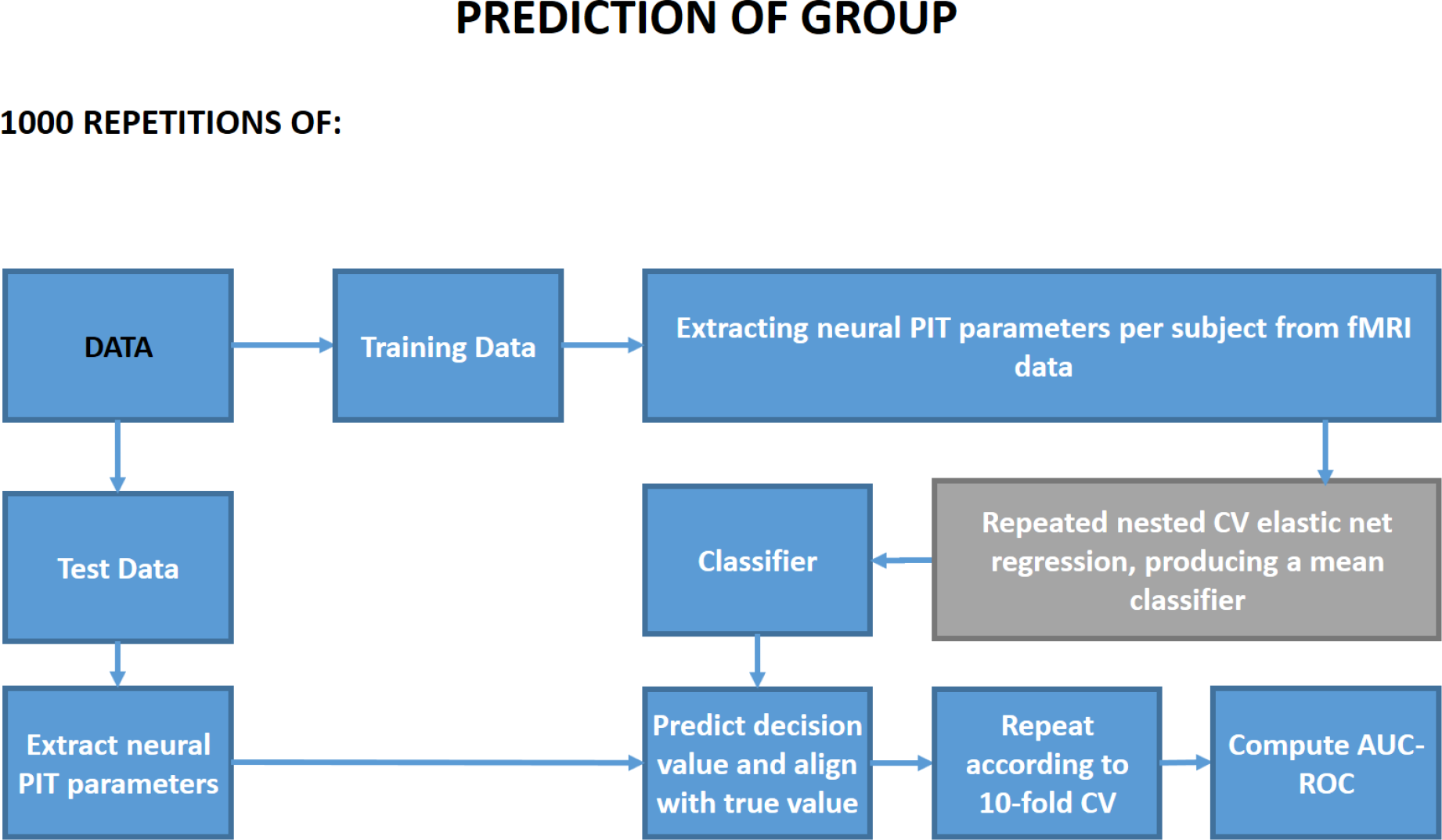
Classifier building algorithm with cross-validation (CV) to estimate generalization error. Nested CV was used for tuning the hyperparameters of the elastic net regression (Varma & Simon, 2006; Zou & Hastie, 2005). This was done repeatedly with different nested CV folds (10 times, 10-fold nested CV) to estimate a robust mean model within each repetition of classifier estimation.

#### Inspecting the classifier based on fMRI data

In order to interpret the final classifier’s regression weights as an *activation pattern* (*a*), i.e. to know how greatly GD and HC differ on all predictor variables we calculated:

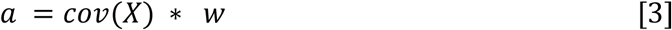

(Haufe et al., 2014), where *w* is the regression weight vector (a column vector), or in other words, the classifier. *X* is the matrix of predictors for all subjects and cov(*X*) is the covariance matrix of *X*.

### Ethics

Subjects gave written informed consent. The study was conducted in accordance with the World Medical Association Declaration of Helsinki and approved by the ethics committee of Charité – Universitätsmedizin Berlin.

## RESULTS

### Cue ratings

Gambling cues were seen as more appropriately representing one or more gambling games than any other cue category: gambling > neutral (β = 1.509, p < 0.001), gambling > negative (β = 1.142, p < 0.001), gambling > positive (β = 1.459, p < 0.001). They elicited more craving in GD subjects than in HC subjects (1.749 compared to 0.719, p < 0.001). Note, that also HC subjects indicated significantly more craving in response to gambling cues compared to neutral cues (β = 0.719, p < 0.001). GD subjects did not rate gambling cues as more positively valenced than HC (β = −0.055, p < 0.712). GD subjects did not rate gambling cues as more arousal-inducing compared to HC subjects (0.142 vs. 0.047, p = 0.525) Also, HC subjects did not rate gambling cues as more arousal inducing than neutral cues (β = 0.047, p = 0.662).

Negatively valenced cues were seen as evoking smaller valence ratings than all other categories in both groups: negative < neutral (β = 0.651, p < 0.001), negative < positive (β = 1.538, p < 0.001), negative < gambling (β = 0.977, p < 0.001). Negative cues were rated similarly by both groups.

Gambling cues lead to higher dominance ratings (β = 0.368, p < 0.001), as did positive cues (β = 0.683, p < 0.001), while negative cues lead to lower dominance ratings (β = -0.297, p < 0.001) in GD and HC subjects compared to neutral cues. GD subjects rated gambling cues as more dominance inducing than HC subjects (β = 0.328, p = 0.021) (**Fig. S2**)

### Describing the behavioral choice data

Here we present results comparing the **laec** model against the **laecg** model (i.e. with an effect of group onto the fixed effects of gain, loss, ed and category). Comparing the two models, we observed a significant χ^2^ difference test result (χ^2^ = 22.6, df = 7, p = 0.002; with ΔAIC = 8.6, ΔBIC = -43.0). Inspecting the estimated parameters of the **laecg** model, we observed that acceptance rate during neutral images with all other parameters at zero (i.e. at their mean, except for *ed*, actually zero) was for HC: 59.3% and for PG: 38.8%, p_Wald_ = 0.119. Gambling cues were associated with stronger increase in gamble acceptance in GD subjects (Δ% = 45) than in HC subjects (Δ% = -8, p_Wald_ = 0.003). The same was true for negative (GD: Δ% = 24, HC: Δ% = -16, p_Wald_ = 0.024) and positive cues (trendwise) (GD: Δ% = 39, HC: Δ% = -2, p_Wald_ = 0.055) (**Fig. 4**). For further behavioral results, please see **Supplements**.

**Figure 4:**
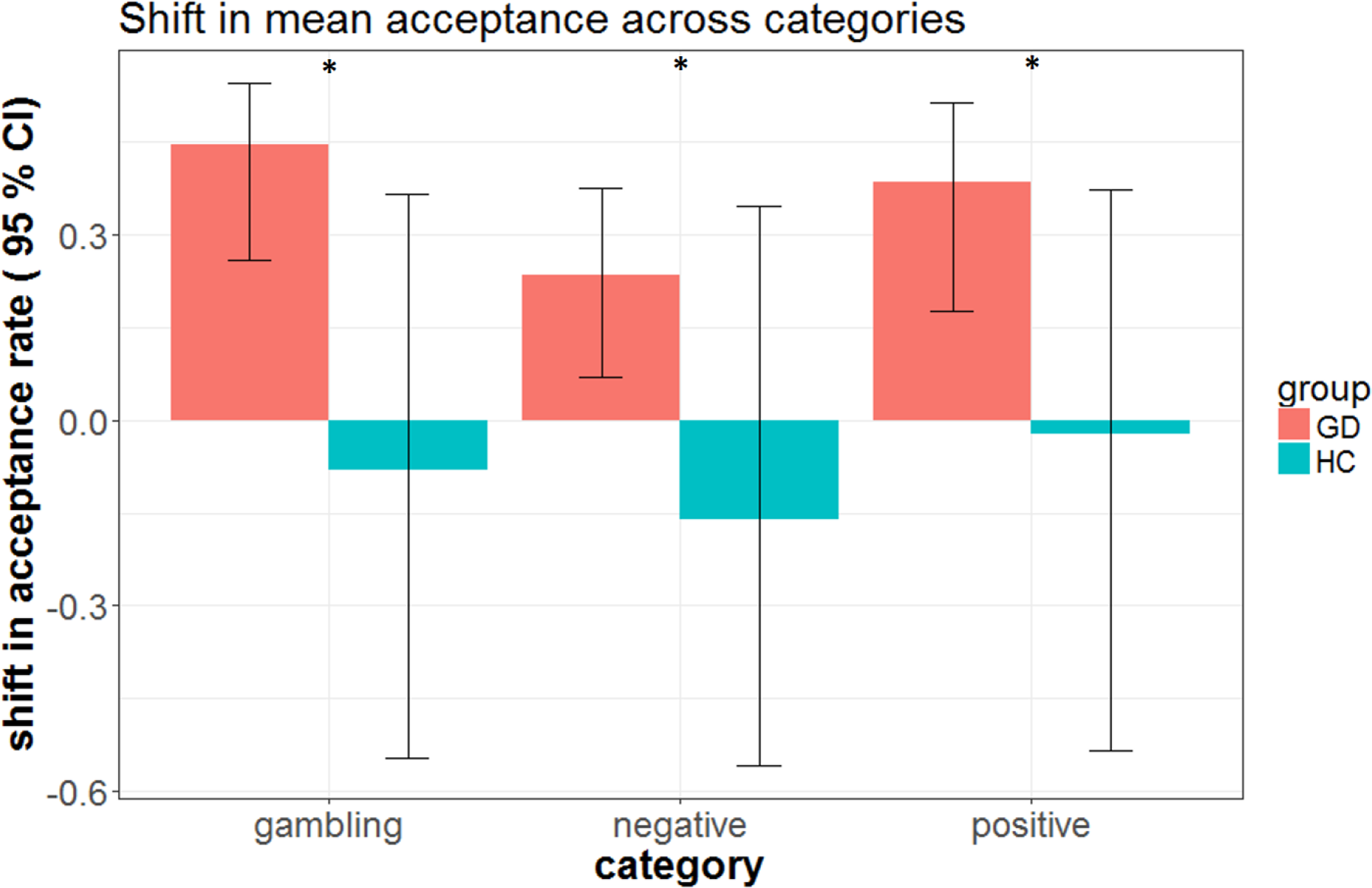
Shift in acceptance rate during gambles per category and group. GD subjects show stronger increase in gamble acceptance (compared to neutral) in comparison to HC subjects during the presentation of all three cue categories in the background.

### Prediction of group using fMRI data

The mean AUC-ROC of the full classifier using neural PIT signatures was 70.0% (mean for the null-classifier, i.e. covariate-only classifier: 61.5%, p = 0.013) (**Fig. 5**).

**Figure 5:**
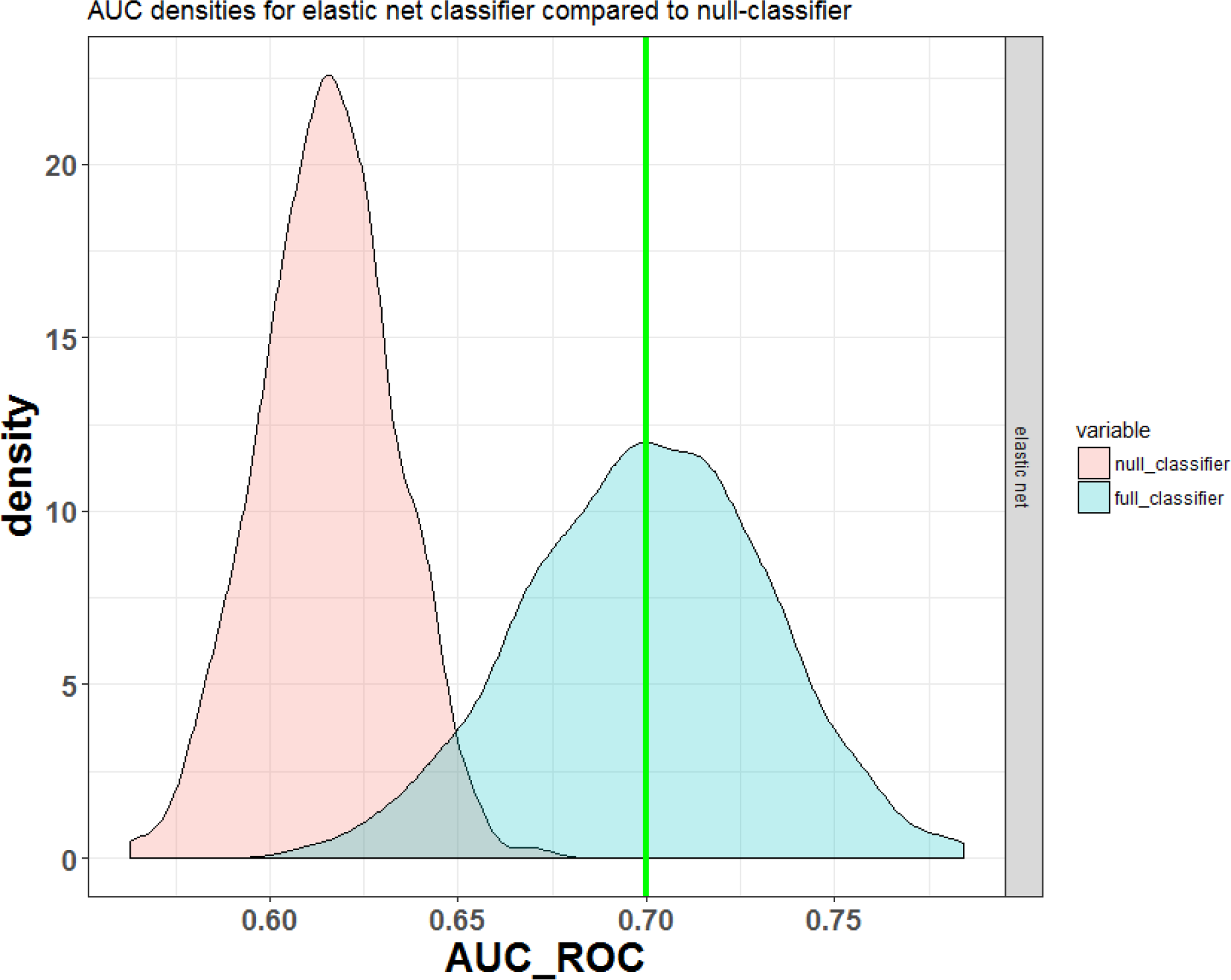
Classification performance of classifier using fMRI neural PIT signatures. Blue is the density plot of 1000 AUC-ROCs obtained from running 1000 repetitions of cross validation of the full classifier using neural PIT signatures. The green line shows the mean of these 1000 AUC-ROCs. In red you see the same density estimate for the null-classifier, i.e. the covariate-only classifier, as a control condition. The full classifier performs significantly better (p_boot_ = 0.013).

We ran the algorithm on the complete data set of fMRI variables. Inspecting the classifier’s logistic regression weights (see **Fig. S3**) (after transformation to activations, see Eq. 3), we saw that the top predictor was negative-cues-PIT-related functional connectivity from amygdala to anterior OFC, with a negative sign. This means that the stronger not accepting a gamble was associated with increase in correlation between amygdala and anterior OFC, the *less* likely the subject was a GD person (and rather a HC subject). In other words, GD subjects showed lower such functional connectivity than HC. The next top three predictors were gambling-cues-related functional connectivity from NAcc to amygdala (positive sign), positive-cues-related functional connectivity from amygdala to lateral OFC (positive sign), and years of education (negative sign) (see **Fig. 6**, **7**).

**Figure 6:**
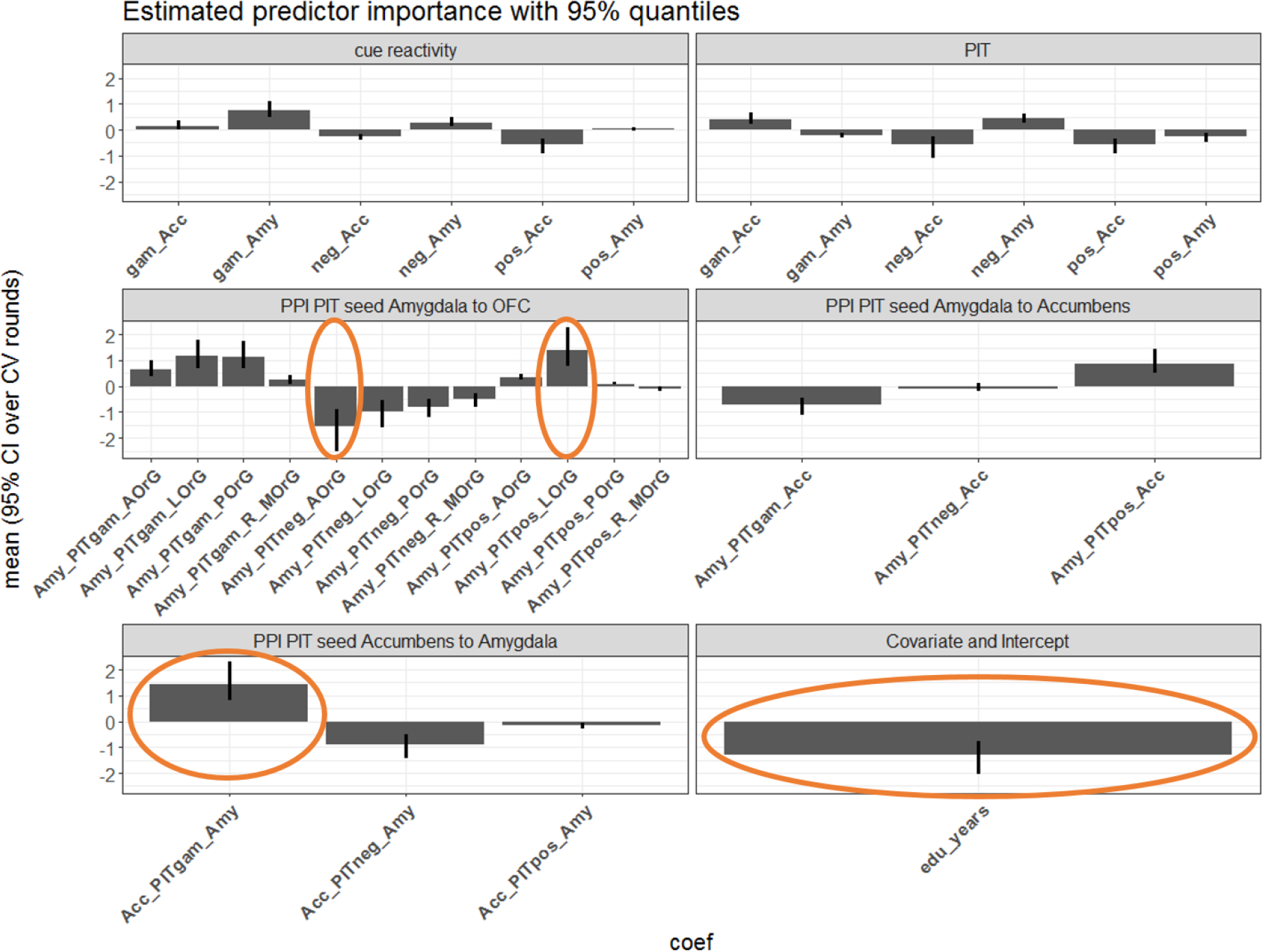
Estimated predictor importance. Plots show estimated predictor importance with 95%-quantiles over 1000 classifier estimation rounds. The larger the absolute size of a importance value the higher the more importantly the difference between HC and GD on the respective predictor maps to classifying a given subject into GD or HC. Positive weights mean GD subjects rather have a higher values compared to HC and negative values vice versa. Red circles mark the regression weights with highest absolute value. Importance values are grouped by the kind of fMRI predictor: cue reactivity related, PIT related, PPI-related. PPIs are further grouped according to seed region and target extraction (e.g. “to OFC”). Icpt: intercept; Acc: Accumbens; PIT: pavlovian-to-instrumental transfer; OFC: orbital frontal cortex; AOrG, LOrG, POrG, MOrG: anterior, lateral, posterior, medial orbital gyrus, i.e. orbital frontal cortex; edu_years: years of education.

**Figure 7:**
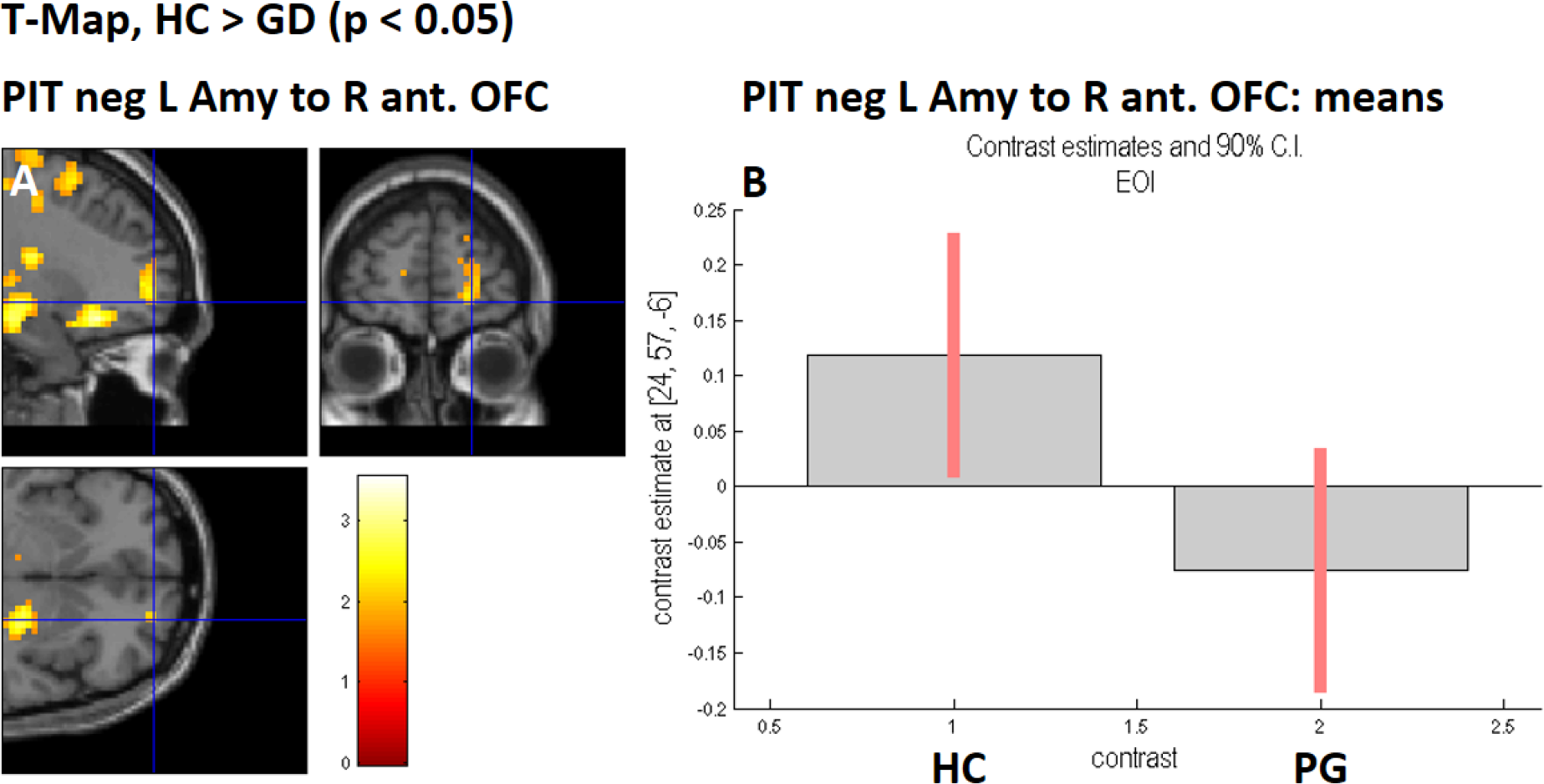
GPPI map for PIT (negative) contrast HC > GD. Displayed at p < 0.05. Illustration of most important weight of the classifier. **A:** from left amygdala to right anterior OFC, centered at peak within region of interest, [24, 57, -6], p_unc_ = 0.024, k = 0. Cluster extends into right superior frontal gyrus. Also visible is significant activity in right posterior OFC, which was also picked up by the classifier. **B:** plot of subjects’ beta values at peak voxel for the contrast in A.

## DISCUSSION

GD is characterized by impaired decision making (Wiehler & Peters, 2015) and craving in response to gambling associated images (Crockford et al., 2005; Goudriaan et al., 2010). There is evidence that the influence of cue-induced emotional states onto value-based decision-making is stronger in GD than HC subjects (Dixon et al., 2006; van Holst, van Holstein, et al., 2012) (Genauck, *under review*). The influence of cues onto value-based decision-making may be regarded as a form of Pavlovian-to-Instrumental Transfer (PIT), the increase of which has been associated with addictive disorders in general (Corbit & Janak, 2007; Garbusow et al., 2016; Genauck, Huys, Heinz, & Rapp, 2013; Mitchell, Gao, Hallett, & Voon, 2016; Schad et al., 2018).

In the current study, we hypothesized that GD subjects should be distinguishable by neural PIT signatures based on fMRI contrasts recorded during an affective mixed-gambles task. We therefore built a classifier using fMRI PIT contrasts to distinguish GD from HC subjects focusing on brain structures known to be relevant in PIT, like amygdala and NAcc (ventral striatum). We also incorporated amygdala’s connectivity to OFC, and amygdala’s and NAcc’s connectivity to each other. We further included neural cue reactivity contrasts as predictors. All these predictors yielded a neural PIT signature per subject which could be used to classify subjects into the GD or HC group.

Our results support our first hypothesis, showing that neural PIT signatures based on fMRI data gathered from the affective mixed-gambles task may successfully classify out-of-sample subjects into GD and HC, with a cross-validated mean AUC-ROC of 70.0% (p = 0.013). This performance on out-of sample data is similar to other studies using MRI data for classification in the field of addictive disorders (Guggenmos et al., 2018; Pariyadath, Stein, & Ross, 2014; Seo et al., 2018, 2015; Whelan et al., 2014). To our knowledge, however, the present study is the first one to use fMRI classification for investigating a behavioral addiction, namely GD, and the neural basis of increased PIT. This means that it is possible to characterize a non-substance related addiction to a considerable degree by a single neuro-functional signature, namely a neural PIT signature in a network of amygdala, NAcc and OFC, derived from PIT and SUD literature. This further implies that addictive disorders, in general, may be associated with PIT-related neural changes, independent of a substance of abuse, which means that neural PIT changes may be a product of addiction-related learning and neural plasticity or even of an innate trait (Barker et al., 2012).

Concerning the predictors in the classifier, we hypothesized that gambling-cue PIT-related functional connectivity from amygdala to OFC should be increased. We found that multiple PIT-related functional connectivities from amygdala to OFC were significant predictors in the classifier. For example, gambling-cues PIT-related functional connectivity from amygdala to OFC was increased in GD compared to HC subjects, in line with our hypothesis and in line with the general prediction that in GD subjects amygdala modulates the value computation in OFC, when addiction-related cues are presented in the background (Cardinal et al., 2002; Holmes et al., 2010; Litt et al., 2008). Furthermore, the top predictor in the classifier was PIT-related functional connectivity from amygdala to anterior OFC in trials with a *negative* cue, with a negative predictor weight. This means that the stronger the rejection of a gamble during the presentation of negative cues was associated with an increase in correlation between amygdala and anterior OFC, the *less* likely the subject was a GD person (and rather a HC subject). In other words, GD subjects showed weaker such functional connectivity than HC. GD subjects, compared to HC subjects, showed significantly more gambling during the presentation of negative cues than during the presentation of neutral cues. HC subjects may not show this effect because of stronger signal transmission related to negative cues from amygdala to OFC. Similarly, it has been found that reduced loss aversion in GD subjects was associated with reduced loss-related functional connectivity from amygdala to ventral medial prefrontal cortex in a pure mixed-gambles task (Genauck et al., 2017). This highlights that impaired decision-making in GD during a pure mixed-gambles task, as well as during an affective mixed-gambles task, may draw from the same functional neural substrate.

We looked at the next two top predictors expecting that PIT-related (as opposed to purely cue reactivity related) neural predictors should be among these. Indeed we found that the next top predictor was gambling-cues PIT-related functional connectivity from NAcc to amygdala (positive sign). This means that the more gamble acceptance during presentation of gambling cues was associated with an increase in correlation between NAcc and amygdala, the *more* likely the subject was a GD person. In other words, GD subjects showed stronger such functional connectivity than HC. NAcc is seen as encoding (especially positive) prediction errors, i.e. it fires when an unexpected reward signal is perceived (Schultz, Dayan, & Montague, 1997). GD subjects rated gambling pictures as more craving-inducing and reacted with significantly stronger gamble acceptance increase than HC when gambling-associated cues were shown in the background. We also saw an important regression weight given to gambling-cues PIT-related functional connectivity from amygdala to OFC, in line with our initial hypothesis. Therefore, it may be that gambling cues elicit a prediction error in NAcc that modulates amygdala activity, which in turn modulates the value representation in OFC in such a way that GD subjects are more inclined than HC subjects to accept the gamble at hand. This is in line with a previous study, where it has been found that GD subjects display increased functional connectivity from amygdala to posterior OFC related to increasing possible gains in a pure mixed-gambles task (Genauck et al., 2017). This highlights again that impaired decision-making in GD during a pure mixed-gambles task, as well as during an affective mixed-gambles task may draw from the same functional neural substrate. Also, it has been observed before that NAcc and amygdala seem to hold relevant signal related to PIT in healthy subjects (Prévost et al., 2012) and to increased PIT in addicted subjects (Garbusow et al., 2016). Interestingly, previous studies (Garbusow et al., 2016; Schad et al., 2018) have observed that in recently detoxified treatment-seeking AD patients, images of alcoholic beverages in the background have a suppressing effect on the instrumental task in the foreground. Contrarily, we have seen that gambling cues elicit a stronger gamble acceptance increase in GD than in HC. This may be because we have included only active non-treatment-seeking gamblers, who at that stage of disease show little activity working against their automated response towards addiction-related cues.

The third top predictor was also PIT related, in line with our hypothesis that PIT-related predictors should be more important than cue reactivity predictors. It was positive-cues PIT-related functional connectivity from amygdala to lateral OFC. This means that the stronger the acceptance of a gamble during the presentation of positive cues was associated with an increase in correlation between amygdala and OFC, the *more* likely the subject was a GD person. In other words, GD subjects showed stronger such functional connectivity than HC. This may be seen as parallel to the finding on behavioral level that GD subjects react with more gambling increase to positive pictures than HC subjects.

Although we discussed the top-three predictors, note that our classifier is truly multivariate. Of the 30 neural PIT signature predictors, 22 received a significant regression weight (and 24 a significant activation weight), despite elastic net regression allowing for total deselection of predictors (Zou & Hastie, 2005). This means that just like on the behavioral level, where GD subjects reacted more strongly than HC to all non-neutral categories, we see that fMRI activities related to *all* non-neutral categories was relevant for characterizing GD. Cue reactivity regression weights are relatively small and the classifier heavily draws on PIT-related variables (the top-three predictors were PIT related). This emphasizes the importance of PIT as a defining marker for addictive disorders beyond cue reactivity.

We used the same cues as Genauck et al. (under review) in a new sample of GD and HC subjects and, in line with that study, we also observed that GD subjects rate the gambling cues as more craving inducing. The ratings and the result that neural PIT signatures successfully distinguish GD from HC subjects reinforce the notion that GD subjects’ cue reactivity facilitates riskier decision-making when addiction-related cues are presented in the background of a gamble task.

Changes in NAcc’s structure (Koehler, Hasselmann, Wüstenberg, Heinz, & Romanczuk-Seiferth, 2015) and function (Koehler et al., 2013; Linnet, Peterson, Doudet, Gjedde, & Møller, 2010; Miedl et al., 2014; Reuter et al., 2005; Romanczuk-Seiferth et al., 2015) related to GD have been observed in previous studies. The same is true for amygdala’s structure (Elman et al., 2012; Takeuchi et al., n.d.) and function (Genauck et al., 2017), as well as for OFC’s structure (Li et al., 2018) and function (Cavedini, Riboldi, Keller, D’Annucci, & Bellodi, 2002; Goudriaan et al., 2010). Our study adds to these findings by considering the functions of these structures concurrently and in a network. Our results support the notion that GD, similar to SUD, is characterized by neural incentive sensitization (Limbrick-Oldfield et al., 2017; Rømer Thomsen et al., 2014) such that in GD a network of amygdala, NAcc and OFC facilitate gambling decisions in the face of gambling cues.

## STRENGTHS AND LIMITATIONS

The main strength of our study is that we have used a classification approach to assess the usefulness of known neural PIT contrasts to characterize GD in out-of sample data. Using this approach, we have estimated the single-subject relevance of these fMRI signals. Our results therefore have not only explanatory value in elucidating the basis of increased PIT in GD, but also predictive value, given that they are likely to be found in new samples of GD and matched HC subjects (Yarkoni & Westfall, 2017). Furthermore, we are to our knowledge the first to address the neural underpinnings of PIT in a behavioral addiction using a machine learning approach. Unfortunately, we have no independent validation sample to externally validate our results (Guggenmos et al., 2018). Further studies are needed to collect such data. As we have laid out, there are multiple ways in which the brain may produce an overt PIT, involving at least amygdala, NAcc and OFC. To increase statistical power, we have omitted other conceptualization of PIT, e.g. as an interference task (Sommer et al., 2017), and hence any limbic-dorso-lateral-prefrontal connectivity (Bray et al., 2008). Future studies should explore this.

## CONCLUSION

We have observed that it is possible to classify HC and GD subjects based on the neural correlates of PIT in a network of NAcc, amygdala and OFC. Our findings further the understanding of GD and show that PIT is relevant for characterizing non-substance-related addictive disorders also on neural level. PIT alterations at the neural level related to an addictive disorder might thus not depend on the direct effect of a substance of abuse, but rather on related learning processes or even on innate traits.

## ACKNOWLEDGMENT

This study was conducted at the BCAN – Berlin Center of Advanced Neuroimaging.

## ONLINE RESOURCES

R code and data (stored in .RData file which is loaded with the R code) to run the classifier estimation and cross-validation, as well as the classical hierarchical regression analyses can be found in the following link. Further you can find there also more detailed data concerning the MRI sequences, as well as the preprocessing of MRI data and the fMRI single subject design: https://github.com/pransito/PIT_GD_MRI_release

