## Supplementary Information for "Neural correlates of cue-induced changes in decision-making distinguish subjects with gambling disorder from healthy controls"

**Title of article:**

**Corresponding Author:**

Alexander Genauck

### Supplements

#### Supplementary methods

##### Administered questionnaires

All subjects completed the Gamblers' Beliefs Questionnaire containing subscales on gambling persistence and gambling illusions (GBQpers, GBQillus) (Steenbergh, Meyers, May, & Whelan, 2002) and the South Oaks Gambling Questionnaire (SOGQ) (Lesieur & Blume, 1987). For matching purposes subjects were asked to indicate age, amount of personal debt and monthly personal net income (Bergh & Kühlhorn, 1994; Ladouceur, Boisvert, Pépin, Loranger, & Sylvain, 1994). They were asked if they were smokers and completed the Fagerström smoking questionnaire (Heatherton, Kozlowski, Frecker, & Fagerström, 1991) (Heatherton, Kozlowski, Frecker, & Fagerström, 1991). Furthermore, they were asked to indicate their level of education which was translated into years spent in primary/secondary school and in tertiary (vocational school/university) education and their handedness. For further characterization of the two groups subjects also completed Beck's Depression Inventory (BDI-II) (Beck, Steer, & Brown, 1996) and the short version of the Barratt Impulsiveness Scale Version 15 (BIS-15) (Patton, Stanford, & Barratt, 1995).

##### Stimuli selection

For the purposes of this study four sets of images were assembled: 1) 67 gambling images, showing a variety of gambling scenes, situations and cues: 36 showing different kinds of slot machines, 12 showing poker, 13 showing roulette, 3 featuring money, 3 featuring dice; 2) 31 images showing negative consequences of gambling (as of now referred to as *negative images*): 7 showing depression / sadness, 4 depicting poverty, 4 depicting debt, 3 showing a quarrel between people, 2 showing family problems, 2 showing the lack of money, 2 symbolizing suicide, 2 showing money burning; 3) 31 images showing positive effects of abstinence from gambling (as of now referred to as *positive images*): 6 showing family, 4 showing relationships, 4 showing friendships, 3 depicting success, 3 depicting freedom, 3 showing joy, 2 showing saved money; 4) 24 neutral images showing objects: 6 kitchen utensils, 8 showing other household objects, 2 showing tools, 2 showing abstract paintings. Important to note, none of the neutral pictures showed humans or faces.

Images were obtained from the internet, sought purposefully to fit the defined categories (positive, gambling, neutral). Online search for images was performed using popular image search engines. Groups of selected images were matched for content as follows: a) percent of images showing a social stimulus (i.e. a person) as opposed to images without persons (gam: 88.2%, pos: 77.4%, neg: 90.3%; $Chi^2 = 1.123$, $df = 2$, $p = 0.570$); b) percent of images showing a face as opposed to people with their face turned away or just hands (gam: 35.3%, pos: 38.7%, neg: 51.6%; $Chi^2 = 3.530$, $df = 2$, $p = 0.171$); c) percent of images showing males (gam: 67.6%, pos: 64.5%, neg: 77.4%; $Chi^2 = 1.300$, $df = 2$, $p = 0.523$).

All images were cropped to fit the aspect ratio optimized to minimize the loss of image area (3:2). Each image was cropped individually making sure that no content was lost. All the images were resized to the resolution of the lowest image in the set (450x300 pixels), ensuring that the image dimensions and quality are the same across all images. Images are available for scientific use upon reasonable request from the first author. They cannot be made publicly available due to copyright.

#### Behavioral analyses

In line with Genauck et al. (*under review*), who used a very similar affective LA task, we used the la model to compare subjects in loss aversion, with…

$$Q = \beta_0 + x_{gain} * \beta_{gain} + x_{loss} * \beta_{loss}$$

Note that one can define $\lambda = -\beta_{loss}/\beta_{gain}$ where $\lambda$ is called loss aversion (Genauck et al., 2017; Kahneman & Tversky, 1979; Tom et al., 2007). Based on the equation defining $Q$ we fit a logistic regression within a generalized linear mixed-effects model, using glmer from the lme4 package in R (Bates, Mächler, Bolker, & Walker, 2015). Here, gain, loss category denoted the fixed effects and subjects and cues and cue category denoted the sources of random effects. To test if the groups differed in the parameters of the **la** model, we expanded the model by an additional fixed effect of group modulating the effect of gain, loss, (**lag**). Statistical testing of the model comparison was performed using $\chi 2$-square difference tests, as well as the comparison of Akaike and Bayesian information criterion (AIC, BIC). For statistical tests of single parameters in the **lag** model, we used Wald z-test as implemented in lme4.

Further, and in line with Genauck et al. (*under review*), we also tested the **lacg** model based on …

$$Q = \beta_0 + x_{gain} * \beta_{gain} + x_{loss} * \beta_{loss} + c^T * \beta_c$$

against the **lacig** model, which assumens separate $\beta_{gain}$and $\beta_{loss}$for each category (Charpentier).

#### Functional Magnetic Resonance Imaging data gathering and preprocessing

Scanning was performed on a 3-Tesla clinical whole-body magnetic resonance tomograph (MR Magnetom Tim Trio, SIEMENS, Erlangen, Germany) equipped with a standard 12-

channel phased-array head coil at the Berlin Center for Advanced Neuroimaging at Charité – Universitätsmedizin Berlin. In the T2*-sensitive Gradient-Echo Echo-Planar Imaging (GE-EPI) sequence used during the affective loss aversion (LA) task, 33 slices covering the whole brain were acquired in descending order (TR=2.0s, 3mm thickness, 25% inter-slice gap, TE: 30ms, flip angle: 78°, in-plane resolution: 64 x 64 pixels, voxel size: 3.0mm x 3.0mm x 3.0mm), using SIEMENS automatic online motion correction. Slices were automatically tilted and aligned with the line from anterior to posterior commissure. Additionally, a T1-weighted 3D structural image for anatomical referencing (Magnetization Prepared Rapid Gradient Echo, MPRAGE, voxel size: 1mm x 1mm x 1mm) and a B0 fieldmap for image distortion correction were recorded. Imaging data were processed with Statistical Parametric Mapping (SPM12, http://www.fil.ion.ucl.ac.uk/spm/, Wellcome Department of Imaging Neuroscience, London, UK) running on MATLAB (version: R2014a, Mathworks, Sherborn, MA, USA). The GE-EPI images of every subject were corrected for differences in slice acquisition time. GE-EPI images were registered to the mean GE-EPI image (motion correction). Fieldmaps were used to unwarp non-linear image distortions caused by B0 inhomogenities (Andersson, Hutton, Ashburner, Turner, & Friston, 2001). The T1 image was co-registered to the unwarped mean GE-EPI image using affine spatial transformation. The T1 image was then segmented into tissue classes and transformed into the Montreal Neurological Institute-standard space (MNI). This process yielded linear and non-linear parameters for the transformation between individual and standard space, which were applied to all unwarped EPI images. Finally, these images were spatially smoothed with an isotropic Gaussian kernel (full-width-at-half maximum 8mm).

#### The fMRI single-subject model

Additionally, the head motion parameters obtained during SPM12 motion correction were entered into the model to account for signal fluctuations caused by the interaction of movement and susceptibility (Morgan, Dawant, Li, & Pickens, 2007). After high pass filtering (cut off frequency = 1/128 Hz) and the elimination of high frequency noise by autoregressive (AR(1)) modeling, the General Linear Model (GLM) was fit to the preprocessed EPIs using a restricted maximum likelihood algorithm.

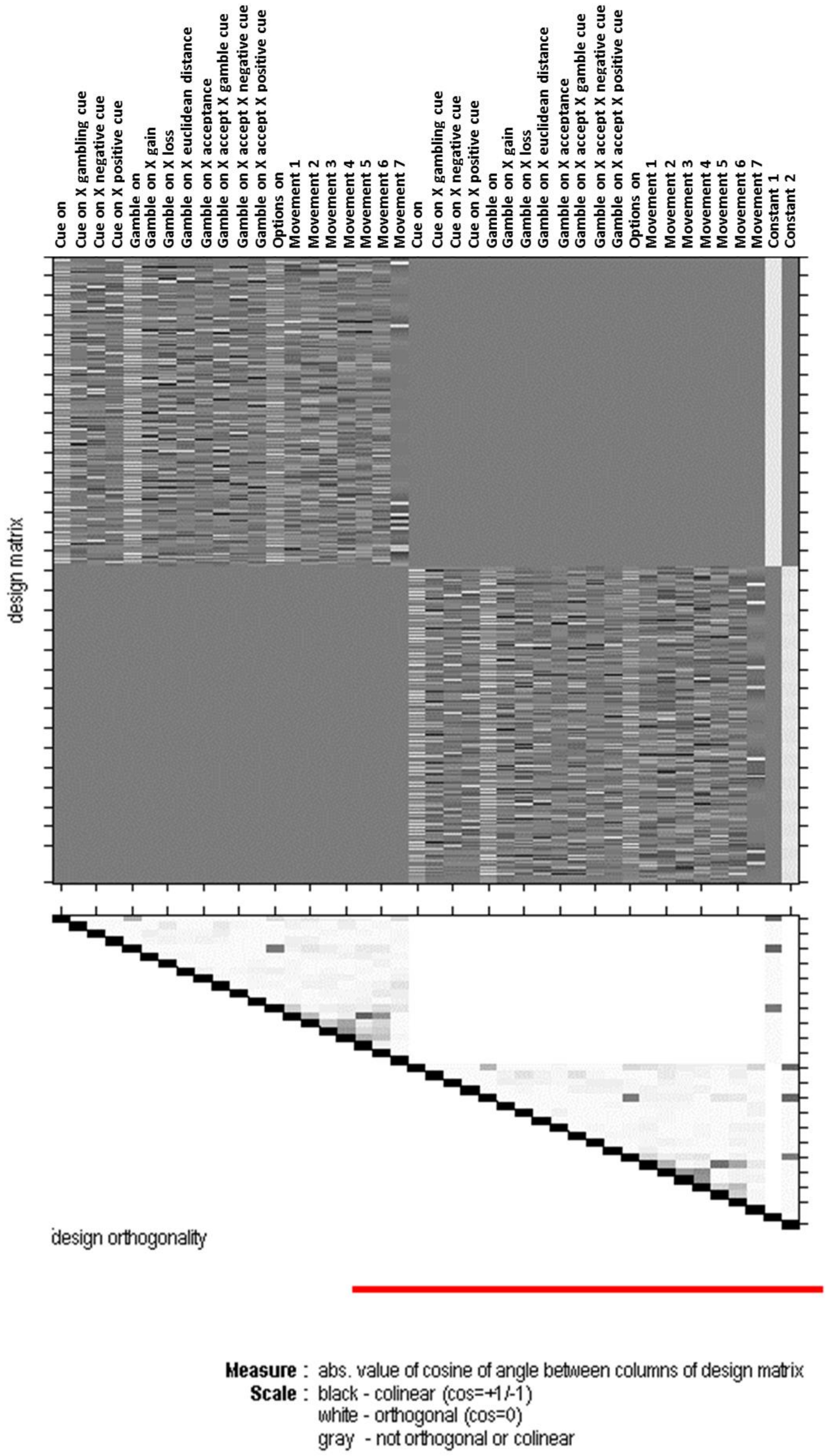


**Figure S1: Design matrix for the single-subject level.** Subjects completed two runs of the affective loss aversion task.

#### Supplementary results

##### Behavioral data

Exploring the **la** model, gain and loss had a significant influence on gamble choice in all subjects ($p < 0.001$, ΔAIC = 4216). There was a fixed effect interaction with group (**lag** vs. **la**) that improved model fit ($p < 0.001$, ΔAIC = 44). In the **lag** model, gain, absolute loss sensitivity, and LA over all trials were for HC 0.30, 0.43, and 1.47, respectively and for GD 0.15, 0.26, and 1.79, respectively. The difference between sensitivity to gain in GD vs. HC was significant ($p_{Wald} = 0.003$). The difference between sensitivity to loss in GD vs. HC was not.

Adding the simple effect of category with group interaction (**lacg**) lead to a significant improvement of the model ($p < 0.001$, ΔAIC = 927). Here, we saw a significantly higher acceptance during gambling ($p_{WaldApprox} < 0.001$), negative ($p_{WaldApprox} = 0.020$), and positive cues ($p_{WaldApprox} = 0.009$) for GD subjects compared to HC. The additional triple-interaction “group X (gain, loss) X category” (**lacig**) did not improve the model ($p = 1$, ΔAIC = -101).

Exploring the parameters of the **laecg** model, during neutral images, sensitivity to gain ($\beta_{gain}$) was for HC 0.27 and for PG 0.14 ($p_{Wald} = 0.003$), sensitivity to loss ($\beta_{loss}$) was for HC -0.39 and for PG -0.23 ($p_{Wald} = 0.059$), and thus loss aversion, based on those fixed effects results, was for HC 1.41 and for GD 1.67.

#### Cue ratings

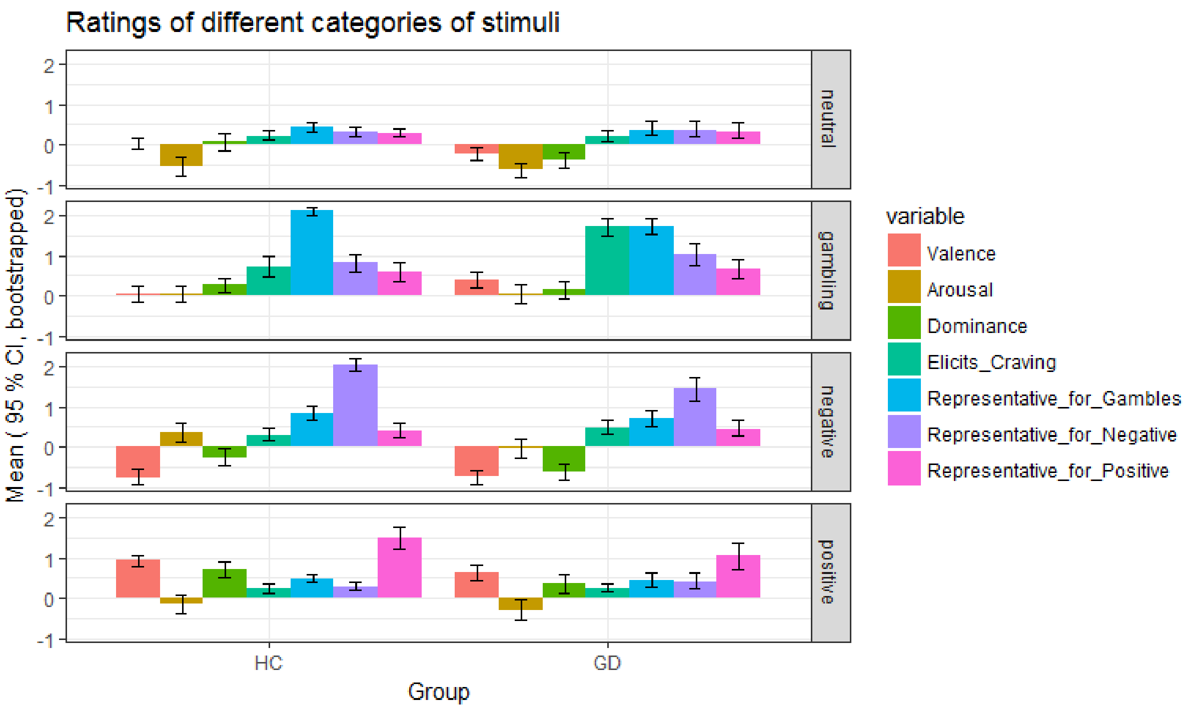


**Figure S2: Means and bootstrapped 95% confidence intervals (CI) of rating variables.** GD: subjects with gambling disorder, HC: healthy controls. Facets report from top to bottom on ratings of neutral category cues, gambling, negative and positive category cues.

#### Regression weights of the fMRI classifier

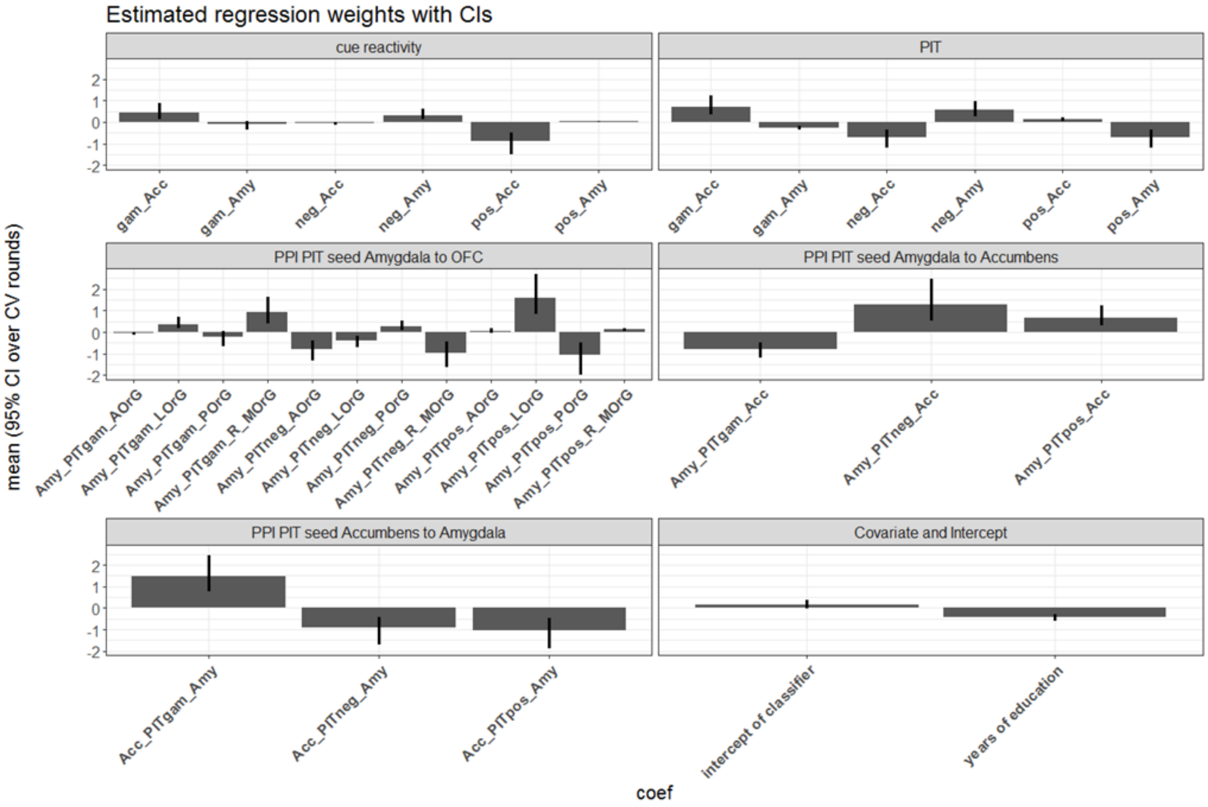


**Figure S3: Estimated regression weights of the classifier when estimated on whole data set.** Plots show regression weights with confidence intervals (95%) across 1000 rounds of classifier estimation. Note that these regression weights cannot be interpreted as predictor importances since they act as a filter which deals with discarding noise and dealing with covariances between variables (Haufe et al., 2014). Regression weights are grouped by the kind of fMRI predictor: cue reactivity related, PIT related, PPI-related. PPI's are further grouped according to seed region and target extraction (e.g. "to OFC"). Icpt: intercept; Acc: Accumbens; PIT: pavlovian-to-instrumental transfer; OFC: orbital frontal cortex; AOrG, LOrG, POrG, MOrG: anterior, lateral, posterior, medial orbital gyrus, i.e. orbital frontal cortex

#### Online materials

R code and data (stored in .RData file which is loaded with the R code) to run the classifier estimation and cross-validation, as well as the classical hierarchical regression analyses can be found in the following link. Further you can find there also more detailed data concerning the MRI sequences, as well as the preprocessing of MRI data and the fMRI single subject design: https://github.com/pransito/PIT_GD_MRI_release
